# Detection and characterization of novel luchacoviruses, genus *Alphacoronavirus*, shed in saliva and feces of meso-carnivores in the northeastern United States

**DOI:** 10.1101/2023.05.31.541188

**Authors:** Ximena A. Olarte-Castillo, Laura Plimpton, Holly McQueary, Yining Sun, Y. Tina Yu, Sarah Cover, Amy N. Richardson, Yuhan Jin, Jennifer K. Grenier, Kevin J. Cummings, Elizabeth Bunting, Maria Diuk-Wasser, David Needle, Krysten Schuler, Michael J. Stanhope, Gary Whittaker, Laura B. Goodman

## Abstract

Small to mid-sized carnivores, or meso-carnivores, comprise a group of diverse mammals, many of which can adapt to anthropogenically disturbed environments. Wild meso-carnivores living in urban areas may get exposed to or spread pathogens to other species, including stray/feral domestic animals. Several coronaviruses (CoVs) have been detected in domesticated and farmed meso-carnivores, but knowledge of CoVs circulating in free-ranging wild meso-carnivores remains limited. In this study, we analyzed 321 samples collected between 2016 and 2022 from 9 species of free-ranging wild meso-carnivores and stray/feral domestic cats in the northeastern United States. Using a pan-CoV PCR, we screened tissues, feces, and saliva, nasal, and rectal swabs. We detected CoV RNA in fecal and saliva samples of animals in four species: fisher (*Pekania pennanti*), bobcat (*Lynx rufus*), red fox (*Vulpes vulpes*), and domestic cat *(Felis catus)*. Next-generation sequencing revealed that all these viruses belonged to the *Luchacovirus* subgenus (*Alphacoronavirus* genus), previously reported only in rodents and lagomorphs (i.e., rabbits). Genetic comparison of the 3’-end of the genome (∼12,000bp) revealed that although the viruses detected group with, and have a genetic organization similar to other luchacoviruses, they are genetically distinct from those from rodents and lagomorphs. Genetic characterization of the spike protein revealed that the meso-carnivore luchacoviruses do not have an S1/S2 cleavage motif but do have highly variable structural loops containing cleavage motifs similar to those identified in certain pathogenic CoVs. This study highlights the importance of characterizing the spike protein of CoVs in wild species for further targeted epidemiologic monitoring.

**Importance:** Several coronaviruses (CoVs) have been detected in domesticated, farmed, and wild meso-carnivores, causing a wide range of diseases, and infecting diverse species, highlighting their important but understudied role in the epidemiology of these viruses. Assessing the viral diversity hosted in wildlife species is essential to understand their significance in the cross-species transmission of CoVs. Our focus here was on CoV discovery in meso-carnivores in the Northeast USA as a potential “hotspot” area with high density of humans and urban wildlife. This study identifies novel alphacoronaviruses circulating in multiple free-ranging wild and domestic species in this area and explores their potential epidemiological importance based on regions of the Spike gene that are relevant for virus-host interactions.

## Introduction

The *Coronaviridae* family encompasses diverse viruses with single-stranded, positive sense RNA genomes. Coronaviruses (CoVs) are currently classified into four genera (*Alpha*-, *Beta*-, *Gamma*, and *Deltacoronavirus*, (1)), that group viruses known to infect birds (mostly gamma- and deltacoronaviruses) and mammals (mostly alpha- and betacoronaviruses), including domesticated and wild animal species, and humans (2). The potential host range of CoVs is coordinated by the spike (S) protein, which is involved in receptor binding via its S1 domain (3). The S protein also mediates membrane fusion with target cells through its S2 domain (4). In general, cleavage of S can occur during virion biosynthesis at a cleavage site between the S1 and S2 domains (S1/S2 cleavage site) which activates the S protein exposing the receptor binding region within the S1 domain, promoting receptor binding (5). After receptor binding, S is also cleaved in the S2 domain close to the fusion peptide (S2’ cleavage site), promoting membrane fusion (6). Coronavirus entry is often regulated through the S cleavage sites. In the *Betacoronavirus* genera, MERS-CoV (from the Merbecovirus clade) can be cleaved by furin at both cleavage sites (S1/S2 and S2’, (5)), while SARS-CoV-2 is the only known member of the Sarbecovirus clade that has a furin cleavage site in S1/S2 (7), which is a key determinant of its emergence (8). In the *Alphacoronavirus* genera, CoVs like feline and canine CoV type 2 (FCoV-2 and CCoV-2, respectively), transmissible gastroenteritis virus and human CoV 229E do not have an identifiable S1/S2 cleavage site (9), while swine acute diarrhea syndrome CoV (SADS-CoV), an emergent and highly pathogenic CoV that possibly originated from bat CoV HKU2 does have an S1/S2 cleavage site (10). The S protein of SADS-CoV also has an additional cleavage site in the S1 domain whose cleavage together with that on the S1/S2 site is essential for cell-cell fusion (11). Overall, multiple cleavage sites may contribute to membrane fusion and virus entry, processes that are crucial determinants for an efficient infection and the disease outcome.

The advancement in sequencing technologies has contributed to the surge of viral discovery which together with comparative genetic analyses can help increase our understanding of how CoV exploit their genetic variation for host adaptation and pathogenesis (3). As the S protein of several CoVs has been crystalized, it is possible to identify and map regions within this protein that interact with the host for receptor binding (12) and cleavage-mediated activation or fusion (13). Comparative genetic analyses of the S protein between CoVs from different hosts can help guide experimental assays to evaluate the importance of these regions and assess the impact of certain mutations on CoV host range and/or pathogenicity. For example, furin, tryspin-like proteases and cathepsins, among other proteases, can cleave the S protein after recognizing certain amino acid sequence motifs (14). Mutations and/or insertions/deletions in these motifs can impact the virus-host interactions. For example, the S1/S2 cleavage site of FCoV type 1 (FCoV-1) and SARS-CoV-2 is located in exposed loops easily accessible to proteases (15–17). Mutations in these loops that disrupt the furin cleavage motif results in highly pathogenic variants for FCoV-1 (18) and variants with decreased pathogenicity for SARS-CoV-2 (19). Likewise, mutations in the S2’ cleavage site can inhibit viral fusion which in MERS-CoV results in reduced viral infectivity (20). Therefore, identifying and mapping in the tertiary structure of S potential cleavage sites in CoVs detected in different species can shed light on the processes CoVs may use to acquire cleavage sites (21) and its relation to host range, pathogenicity and evolution of CoVs (22).

*Luchacovirus* is a subgenus of *Alphacoronavirus* (also known as rodent or murine *Alphacoronavirus*), that includes viruses infecting rodents (order Rodentia) and rabbits (order Lagomorpha). Although currently classified as alphacoronaviruses, the S gene of luchacoviruses is more similar to and groups with betacoronaviruses and not alphacoronaviruses, likely as a result of an early recombination event (23). To date, luchacoviruses have only been detected in at least 13 species of rodents (including rats, mice, and voles) and 2 species of lagomorphs (including rabbits and plateau pika, Table 1). Even though luchacoviruses are highly diverse and have been reported from various rabbit and rodent species in different countries in Asia, Europe and the U.S. (24) (25) they form a monophyletic group in the phylogenies involving the S and other genes and share similar genetic features in their genomes, suggesting a long association with rodents and rabbits (23). In general, the S gene of these viruses is highly variable and the sequence of one of these viruses (JC34, accession number KX964649) obtained from a Chevrier’s field mouse (*Apodemus chevrieri*) in China in 2011 (26) has a furin cleavage motif (S-R-R-A-R | A, where S is Serine, R is Arginine, A is Alanine and | is where the cleavage occurs) in its S1 domain. Interestingly this motif has a very similar amino acid sequence to the S1/S2 cleavage site of SARS-CoV-2 (P/H-R-R-A-R | S, where P is proline, H is Histidine (22)), which is an atypical furin recognition motif (the canonical furin cleavage motif is X-R-R-X-K/R-R | S, where K is lysine and X is any residue). Although experimentally the luchacovirus JC34 cleavage motif is not cleaved by human furin (27), continued surveillance of these viruses focusing on targeted regions is essential to understand if there is variation in regions with motifs of interest and their relevance in the origin and emergence of new CoVs.

**Table 1.**
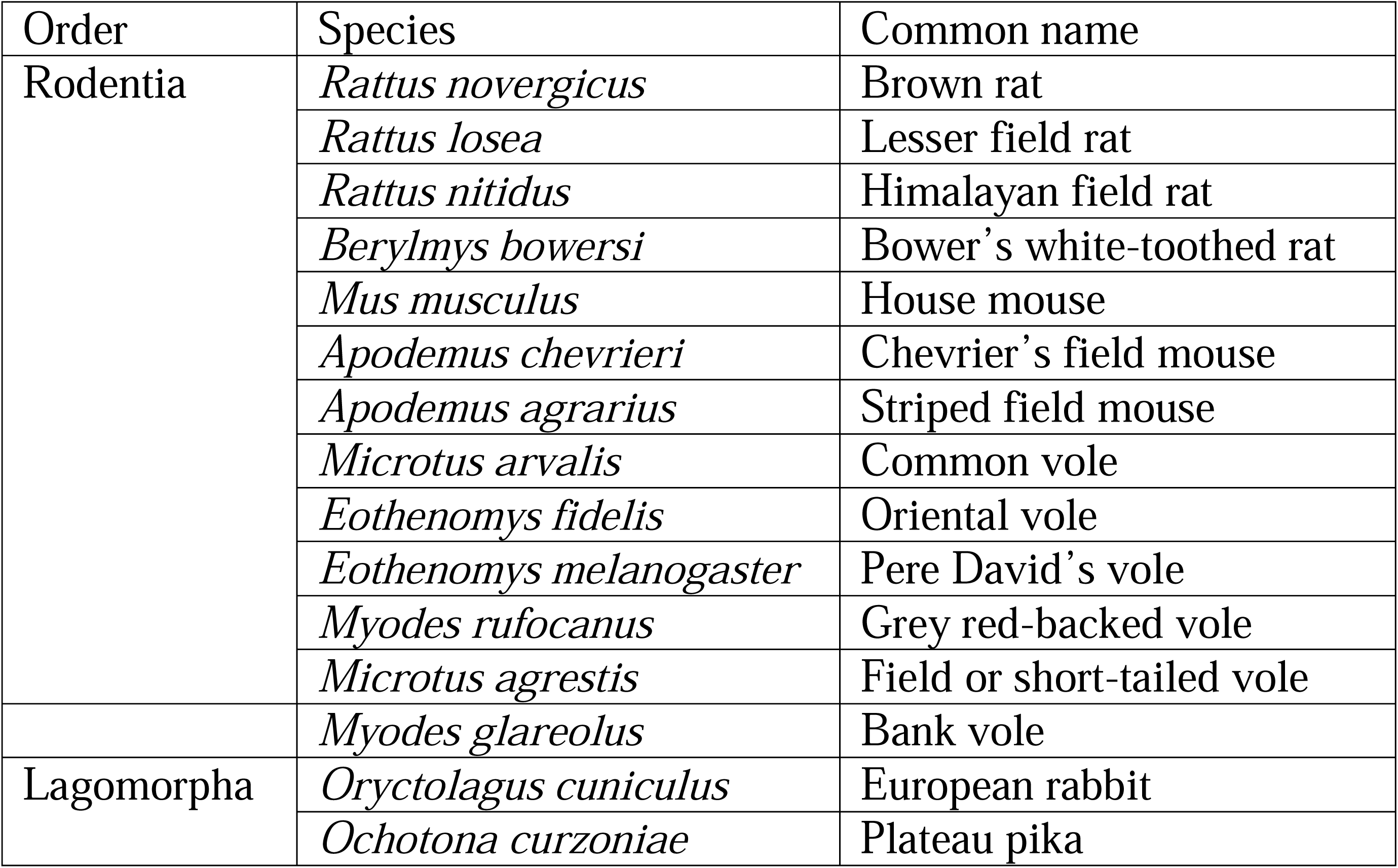
Species of rodents and lagomorphs from which Luchacoviruses have been reported.

The meso-carnivores are a group of small to mid-size carnivores of economic and ecologic relevance that include domesticated and farm animals (dogs, cats, and American mink, *Canis lupus familiaris, Felis catus, Mustela vison,* respectively) as well as various wild species. In the northeastern United States (U.S.), there are 14 species of meso-carnivores (28), including ubiquitous species like raccoons (*Procyon lotor*), red foxes (*Vulpes vulpes*), and striped skunks (*Mephitis mephitis*). Due to their diverse behavior and ecology, meso-carnivores can thrive in various habitats including urban and rural areas (29). In rural areas, they may act as reservoirs for pathogens that may spill over into other species of wild carnivores. Indeed, skunks and raccoons in the eastern U.S. can maintain the sylvatic cycle of rabies independently of domesticated animals (30). In urban areas they may get exposed to pathogens from or spread infectious diseases to domesticated species, especially free-ranging stray or feral dogs and cats. In the U.S., early estimates suggest there may be more than 30 million free-ranging feral/stray cats (31). Feral/stray cats often live in large groups (known as feral cat colonies) in which some viruses are highly prevalent (32)and large-scale outbreaks can occur (33). In many cities in the U.S., including New York City (NYC), humans often provide food for stray cats through feeding stations. These stations also attract wild species, facilitating the interspecific contact between domestic cats and other urban meso-carnivores as well as with rodents. Despite their potential importance in the spread of infectious diseases, our knowledge of the viral diversity hosted by free-ranging wild and domesticated meso-carnivore species in the U.S. remains limited (34).

Alpha- and betacoronaviruses can infect diverse species of meso-carnivores causing a range of diseases. Alphacoronaviruses like mink CoV (35) and ferret enteric CoV (36) cause enteric diseases in mink and ferret, respectively, while ferret systemic CoV (36) and FCoV (37) can cause highly fatal diseases in ferrets and domesticated cats, respectively. Betacoronaviruses like canine respiratory CoV are widely spread throughout the U.S. causing respiratory signs in domesticated dogs (38). Like their domesticated counterpart, wild meso-carnivores also host different CoVs, including the divergent novel alphacoronaviruses identified in free-ranging Asian leopard cats (*Prionailurus bengalensis*) and Chinese ferret badgers (*Melogale moschata*) in southern China (39). Meso-carnivores also play a role in the epidemiology of CoVs relevant to humans including SARS-CoV-2 which can infect minks and ferrets (40), species that can also infect humans (41). Likewise, recent studies have found CCoV-2, a common *Alphacoronavirus* of domesticated dogs and wild canids, in humans (42). This zoonotic event was related to extensive recombination with other alphacoronaviruses and the possible loss of function of the N-terminal region of the S1 domain (43). Meso-carnivores may also be susceptible to CoVs from domesticated and farmed meso-carnivores, due to their close genetic relatedness. For example, the sequence of the receptor for CCoV-2 and FCoV-2 (Aminopeptidase N) is highly similar (>92.7%) between members of the Felidae and Canidae families (44). Overall, both urban and rural wild meso-carnivore carry a high diversity of viruses (45) that need to be further explored to understand the complex epidemiology of CoVs in wild populations, especially those highly adapted to anthropogenically disturbed areas.

The aim of this study was to perform a longitudinal surveillance of CoV in free-ranging wild meso-carnivore in the northwestern region of the U.S. Using next generation sequencing, comparative genetic and phylogenetic analyses, we provide the first molecular characterization, focusing on the S gene, of luchacoviruses detected in three species of meso-carnivore in the U.S. and show that the luchacoviruses detected in the U.S. to date, are highly diverse when compared to those reported elsewhere.

## Methods

### Sample collection

Between 2016 and 2022, tissues and feces from 9 species of meso-carnivores including bobcat, fisher, American mink, coyote, red fox, racoon, striped skunk, river otter (*Lontra canadensis*), and gray fox (*Urocyon cinereoargenteus*) were collected. The samples were opportunistically collected from carcasses turned in for routine necropsy to either the New York State or New Hampshire veterinary diagnostic laboratories. In 2021 samples from 4 species of rodents including chipmunk (subfamily Xerinae), Eastern cottontail (*Sylvilagus floridanus*), Eastern gray squirrel (*Sciurus carolinensis*), and North American porcupine (*Erethizon dorsatum*) were also collected.

In NYC 30 rectal swabs from 17 raccoons and 7 striped skunks between August and October 2021, and rectal, saliva, and nasal swabs from 7 feral cats in August and September 2022 were collected. Animals were captured in Tomahawk Live Traps baited with cat food. Traps were baited and set at dusk and checked at dawn. Captured animals were anesthetized using ketamine-dexmedetomidine administered by a veterinarian in the field. Each sedated animal was given a routine health exam and rectal swabs were collected from each individual using sterile polyester tipped swabs by gently inserting the swab into the anus and rotating the rod on its axis for 10-15 seconds. Swabs were then individually stored in 1.5mL screw top microcentrifuge tubes with 500uL of RNAlater (Thermo Fisher Scientific). Samples were transported on ice, incubated at 4°C overnight and transferred to -80°C for long term storage prior to RNA extraction. After sample collection, animals were returned to their cage, monitored during recovery, and released at the point of capture. Capture and sampling of NYC meso-carnivore was performed under New York State Department of Environmental Conservation (NYSDEC) permit #2704 and were approved under Columbia University’s IACUC protocols #AC-AABA1469 and #AC-AABA1472

### Pancoronavirus screening

Total RNA was obtained from tissues, feces, and swabs using the MagMax *mir*Vana Total RNA isolation kit (Thermo Fisher Scientific) using the KingFisher Apex Purification System (Thermo Fisher Scientific). Prior to RNA extraction between 20 to 40 mg of sample was disrupted using the Bead Ruptor Elite (OMNI International) and 800 ul of lysis buffer. For the disruption, MagMAX Microbiome Bead Tubes were used for feces or rectal swabs and 500mg of ∼2.5mm Zirconia/Silica beads (Biospec Products) were used for tissues and saliva and nasal swabs. The obtained RNA was reverse transcribed using the SuperScript VILO Master Mix (Thermo Fisher Scientific). CoV RNA was detected using a pancoronavirus nested PCR with primers targeting the conserved RdRp gene of CoVs as previously published (Watanabe et al., 2010) and using the Platinum II Taq Hot-Start DNA Polymerase (Thermo Fisher Scientific). After PCR, samples were run in a 1% agarose gel stained with GelRed (Biotium) and visualized in a Gel Doc XR System (Biorad). Positive samples were Sanger sequenced at the Biotechnology Resource Center (BRC) at Cornell University.

Confirmed CoV-positive samples were further sequenced by RNA-seq with a sequencing depth of 100 million reads at the Transcriptional Regulation and Expression Facility at Cornell (RRID:SCR_022532). RNA-seq libraries were prepared using the NEBNext Ultra II Directional RNA Library Prep Kit for Illumina (New England Biolabs, NEB) using 200-600ng total RNA as input. Subtraction of both bacterial and mammalian rRNA was done using the NEBNext rRNA Depletion Kits (NEB), and sequenced on a Novaseq 6000 instrument (Novogene). RNA Reads generated were uploaded to the Chan Zuckerberg ID (www.czid.org, (46)) to find matches to nucleotide and protein sequences available in NCBI. To close gaps from the sequences obtained after RNAseq, we designed primers based on the known sequences and performed PCR on the previously obtained cDNA using Platinum II Taq Hot-Start DNA Polymerase (Thermo Fisher Scientific). Obtained products were sequenced in a MinION Mk1C (Oxford Nanopore Technologies, ONT) using a Flow Cell R10.4 (ONT) and the Native Barcoding Kit 24 V14 (ONT).

### Genetic and phylogenetic analyses

In this study the partial sequences of the RdRp gene (401nt) for five positive samples (W291/bobcat/2021, W145/fisher/2022, W317/red fox/2022, DC6837 F/domestic cat/2022 and DC6837 S/domestic cat/2022), the partial sequence of the S gene (2,561nt), that includes the complete S1 domain and a partial sequence of the S2 domain, of one of our positive samples (W291/bobcat/2021) and the complete 3’-end of the genome (12,214nt) of one of our positive samples (W317/red fox/2022) were obtained. These seven sequences were deposited in GenBank (accession numbers OQ756331-35 and OR428266-7). The five partial sequences of the RdRp (401nt) and the two complete sequences of the S1 domain of the S gene (1674nt) obtained in this study were each aligned with homologous sequences from 60 and 36 other CoVs from the four families, respectively. These alignments were obtained using the Clustal Omega algorithm (47) and each was used to compute the best-fitting nucleotide substitution model and to construct a maximum likelihood (ML) tree using MEGA 11 (48).

Open reading frames (ORFs) were detected in the complete sequence of the 3’-end of the genome (12,214nt) obtained from one of our samples and 15 other luchacoviruses sequences available using Geneious Prime 2023.0 (Dotmatics). These 16 sequences were aligned using the MUSCLE algorithm (49) after which we deleted the regions where gaps were identified and then generated a similarity plot using SimPlot 5.1 (50). This alignment was also used to find possible recombination breaking points using the Recombination Detection Program (RDP) 5 (51) using the RDP method (52). Additionally, the partial S gene sequence obtained from one of our samples (2,561nt) was aligned with the 16 luchacovirus sequences mentioned previously using the MUSCLE algorithm (49). This alignment was used to generate a similarity plot using SimPlot 5.1 (50). These 17 partial S sequences were translated into their respective amino acid sequences and aligned with the homologous amino acid sequences of two alphacoronaviruses, SADS-CoV (MT294722), bat CoVs HKU2 (NC_009988), and three betacoronaviruses including SARS-CoV-2 (QQN72495), a rodent CoV (MT820632) and a rabbit CoV (JN874560) using the Clustal Omega algorithm (47). This alignment was used to compare regions in S in which known cleavage sites occur including the S1/S2 and S2’ regions of SARS-CoV-2 (53), and regions in the S1 domain in which *Luchacovirus* JC34 (22) and SADS-CoV (11) have predicted and characterized furin cleavage sites, respectively. We detected potential cathepsin cleavage sites by looking for specific amino acid motifs previously identified for human cathepsins L, V, K, S, F, B (14). Furin cleavage sites were predicted using ProP (54) (ProP values >0.5). Regions of interest (i..e. possible furin cleavage sites) were mapped on the tertiary structure of the S protein of SADS-CoV, which is the most closely related *Alphacoronavirus* to the *Luchacovirus* clade that has an available crystalized tertiary structure (55). The colors of the protein and the mapping of the regions of interest were done using Chimera X (56).

## Results

From 184 individual meso-carnivores we collected 321 samples: 152 feces (or rectal swabs), 7 saliva swabs, 7 nasal swabs, and 155 tissues samples (Table 2) including liver (43%), lung (44%), brain (2.8%), kidney (6%), small intestine (2.8%), and heart (1.4%) from 9 species (Table 3). Using a nested PCR pancoronavirus screening method we found 5 samples positive for CoV RNA, 4 from feces and 1 from a saliva swab (Table 2) from three different meso-carnivore species (Table 3). The species with positive samples included a bobcat collected in October 2021 (W291/bobcat/2021), a fisher collected in January 2022 (W145/fisher/2022), a red fox (W317/red fox/2022) collected in February 2022, and a feral domestic cat collected in August 2022 from which we found a positive sample in the saliva swab (DC6837 S/domestic cat/2022), and in the rectal swab (DC6837 F/domestic cat/2022), Table 4). The bobcat and red fox samples were collected in New Hampshire and the fisher and feral domestic cat samples were collected in New York State. Additional tissues or swabs from the four animals that were positive for CoV were screened and all were negative for the presence of CoV RNA. The other tissues included the lung from the bobcat (W291/bobcat/2021) and red fox (W317/red fox/2022), the lung, liver, kidney, and heart from the fisher (W145/fisher/2022), and the nasal swab from the feral cat (DC6837 F and S/domestic cat/2022). Of the 30 fecal samples from raccoons and skunks sampled in 2021 in New York City, none was positive for any CoV. For rodents, we collected 36 samples in 2021, including feces (50%) and lung (50%). We did not find CoV in any of these samples (Table S1).

**Table 2.** Results of the CoV screening of meso-carnivore according to sample type.

| Sample type | Total | Positive | Negative |
| --- | --- | --- | --- |
| Tissues | 155 | 0 | 155 |
| Feces or rectal swabs | 152 | 4 | 148 |
| Saliva swabs | 7 | 1 | 6 |
| Nasal swabs | 7 | 0 | 7 |
| Total | 321 | 5 | 316 |

**Table 3.** Results of the CoV screening of meso-carnivore according to species.

| Species | Total | Positive | Negative |
| --- | --- | --- | --- |
| Fisher | 124 | 1 | 123 |
| Bobcat | 40 | 1 | 39 |
| Raccoon | 39 | 0 | 39 |
| Coyote | 24 | 0 | 24 |
| Otter | 21 | 0 | 21 |
| Red fox | 21 | 1 | 20 |
| Domestic cat | 21 | 2 | 19 |
| Skunk | 18 | 0 | 18 |
| Gray fox | 8 | 0 | 8 |
| Mink | 5 | 0 | 5 |
| Total | 321 | 5 | 316 |

**Table 4.** Results of the CoV screening of meso-carnivore according to year of collection.

| Year of collection | Total | Positive | Negative |
| --- | --- | --- | --- |
| 2016 | 1 | 0 | 1 |
| 2017 | 2 | 0 | 2 |
| 2018 | 47 | 0 | 47 |
| 2019 | 71 | 0 | 71 |
| 2020 | 18 | 0 | 18 |
| 2021 | 123 | 1 | 122 |
| 2022 | 59 | 4 | 55 |
| Total | 321 | 5 | 316 |

For the five positive samples obtained from the meso-carnivores, we initially sequenced a partial region of the RdRp gene (401bp). The percentage sequence similarity of this region between these three sequences was 82.3%. The CoV sequences obtained in the feral cat in NYC (DC6837 F and S/domestic cat/2022) were identical to the one obtained from the red fox (W317/red fox/2022). A phylogeny including these five sequences plus 60 others from CoVs in the four genera revealed that all sequences obtained in this study are within the Luchacovirus clade (Figure 1). Sequences W145/fisher/202, W317/red fox/2022, and DC6837 F and S/domestic cat/2022 clustered in a group together with viruses from rabbit and vole reported in Europe, while W291/Bobcat/2021 did not group with any specific group of luchacoviruses. The Luchacovirus cluster grouped together with SADS-CoV and bat CoV HKU2 (Figure 1, emphasized with blue arrows).

**Figure 1.**
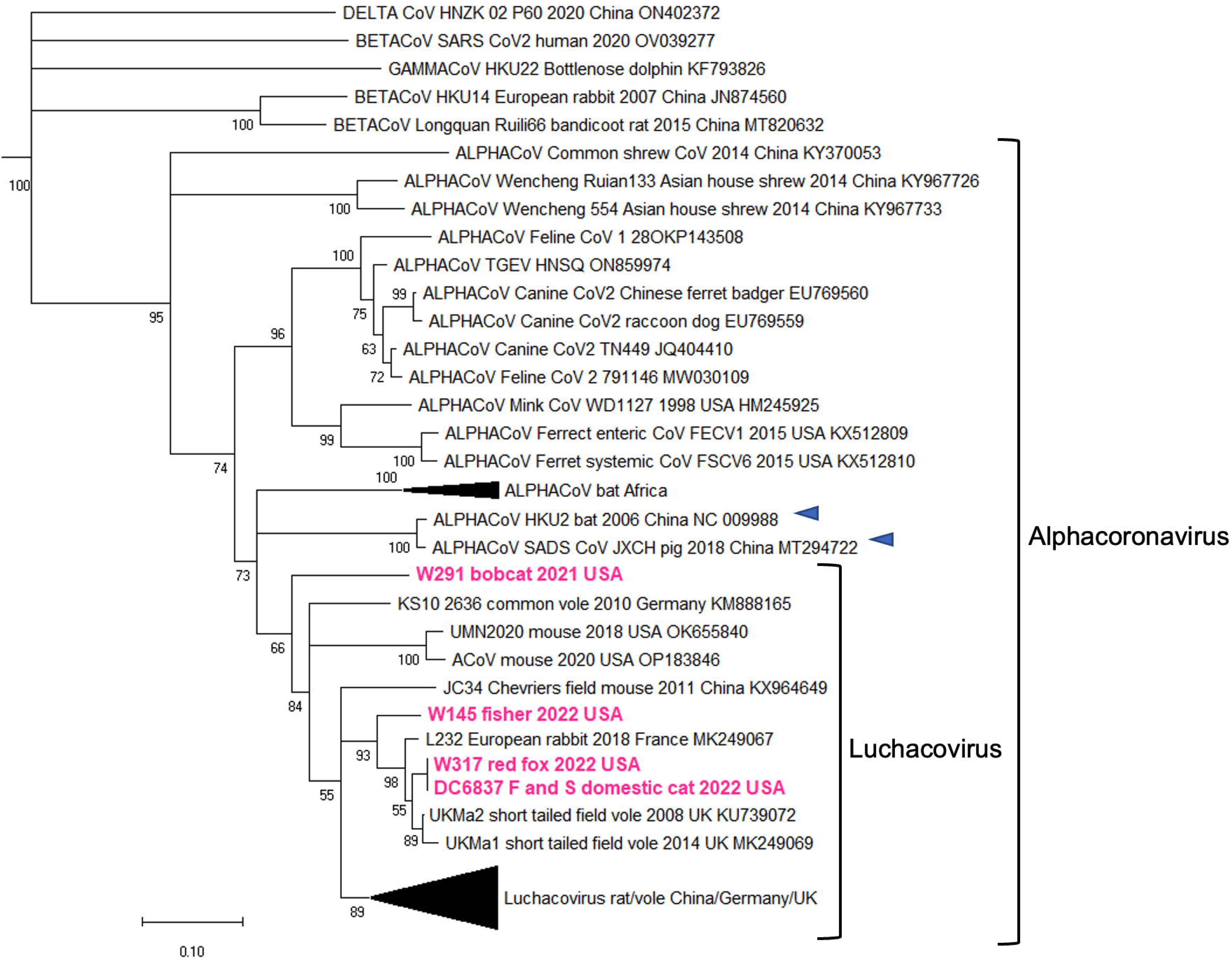
Maximum likelihood phylogenetic tree of a partial region of the RdRp gene of 65 coronaviruses (CoVs) from the four genera, including five obtained in this study (highlighted in fuchsia). The Luchacovirus clade is within the *Alphacoronavirus* group, each shown within labelled brackets. The CoVs most closely related to the Luchacovirus group (SADS-CoV and bat CoV HKU2) are emphasized with blue arrows. For all viruses the virus name, host species, year, and country of collection, and Genebank accession numbers are shown at the tip of the tree. Numbers at the branches indicate bootstrap percentage values from 1000 replicates. Branches with support <50 were collapsed. Nucleotide substitution model used: HKY+I+G.

Additional sequences were obtained for W317/red fox/2022 and W291/bobcat/2021 but not for W145/fisher/2022 or DC6837 F and S/domestic cat/2022 due to low quality and quantity of RNA. For W317/red fox/2022 we obtained a partial genome sequence (12,214nt) that includes the complete S gene and for W291/Bobcat/2021we obtained a partial sequence of S gene (2,561nt) containing the complete sequence of the S1 domain and a partial sequence of the S2 domain. A phylogeny constructed using the nucleotide sequences of the S1 domain of the two variants obtained in this study and 36 other CoVs from the four families revealed that our sequences also grouped within the Luchacovirus cluster and that this group was not closely related to the *Alphacoronavirus* group (Figure 2). In general, there were three main groups (bootstrap support of 100%) within the Luchacovirus cluster: Group 1 includes viruses from voles and rat from China, Group 2 includes viruses from rat from China and the UK and Group 3 includes viruses from mice and rat from China (Figure 2). There were 4 luchacoviruses, including the two reported in this study, that did not cluster within these three groups. W317/red fox/2022 was placed in a group with one of the other divergent luchacoviruses obtained from a pika (P83/plateau pika/China), outside Group 3 while W291/Bobcat/2021clustered outside groups 1 and 2. The fourth divergent luchacovirus, UMN2020/mouse/2018/USA, collected from a mouse in the U.S., did not group with any other luchacovirus (Figure 2). Each of the three defined groups has an overall higher percentage sequence similarity in the S1 domain (>92%) than when compared to the other groups (group 1 vs 2 50%, group 1 vs 3 56.2% and group 2 vs 3 51.8%) or to the 4 luchacoviruses that were not within these groups, including the two viruses obtained in this report, W317/red fox/2022 and W291/Bobcat/2021 (<64%), as well as P83/plateau pika/China (<69%) and the most divergent one, UMN2020/mouse/2018/USA (<52%). Like in the phylogeny of the partial RdRp gene, the Luchacovirus cluster grouped together with SADS-CoV and bat CoV HKU2 (Figure 2, emphasized with blue arrows). Phylogenetic trees of the E, M, and N genes (Figure S1) also show the three well-supported groups within the Luchacovirus clade. In these trees, W317/red fox/2022 did not cluster with any of these groups and only in the tree of the M gene did it cluster with UMN2020/mouse/2018/USA, the luchacovirus detected in a mouse in the U.S.

**Figure 2.**
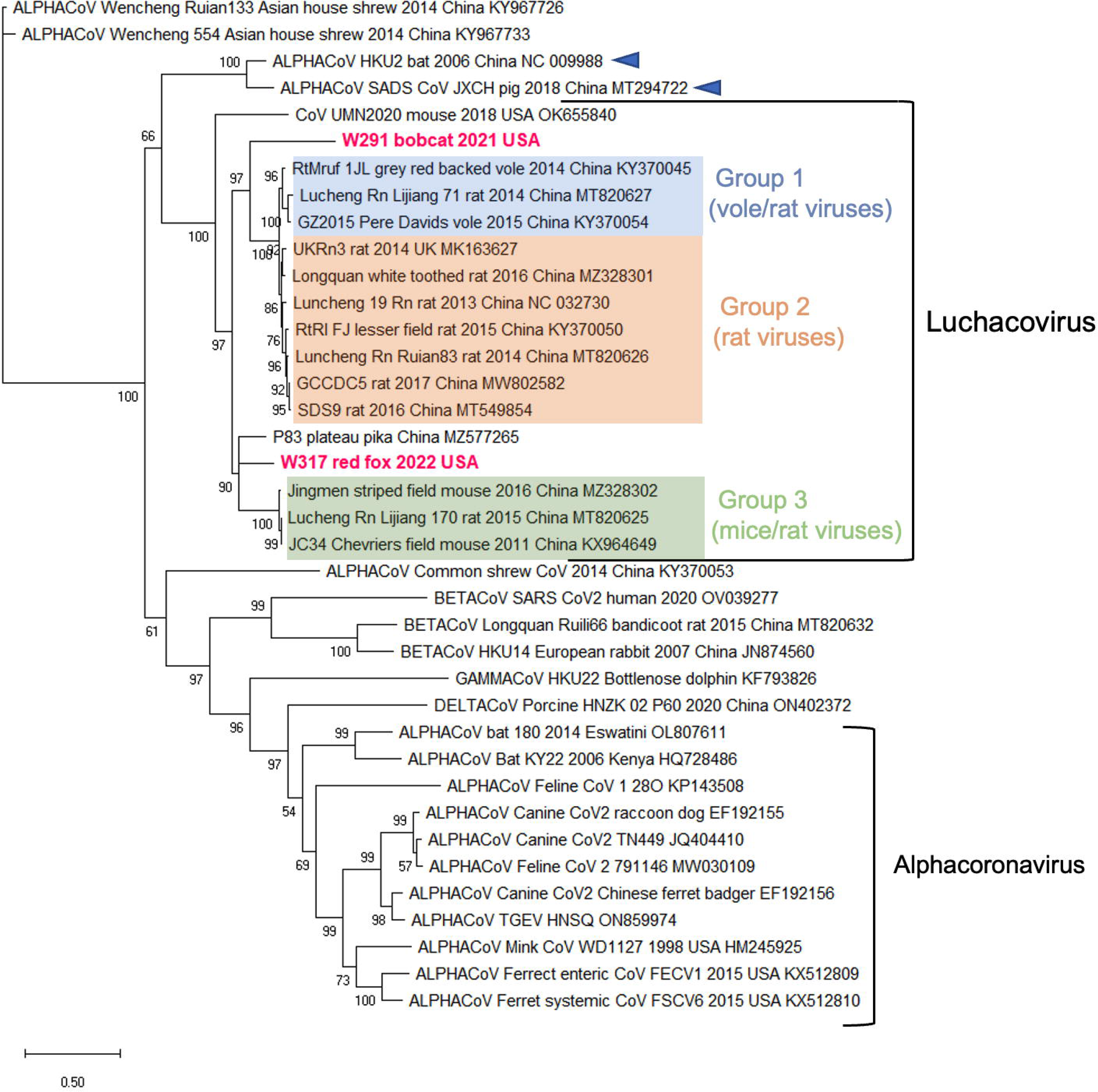
Maximum likelihood tree of the complete nucleotide sequence of the S1 domain of the S gene of 38 coronaviruses from the four genera including the two obtained in this study (highlighted in fuchsia). In this phylogeny the Luchacovirus clade is not within the *Alphacoronavirus* group, each shown within labeled brackets. Within the Luchacovirus group the three well-supported groups (bootstrap of 100%) are shown in different colors (Group 1 in blue, Group 2 in orange, Group 3 in green). Four luchacoviruses, including the ones identified in this study do not cluster within these three groups. For all viruses the virus name, host species, year, and place of collection, and Genebank accession number are shown at the tip of the tree. Numbers at the branches indicate bootstrap percentage values from 1000 replicates. Branches with bootstrap support of <50 were collapsed. Nucleotide substitution model used: GTR+I+G.

The amino acid sequences of the partial S protein (853 amino acids), containing the complete S1 domain and a partial region of the S2 domain of W317/red fox/2022 and W291/bobcat/2021 were 52.1% similar. We compared the amino acid sequences of the four regions in S known in other CoVs to have protease cleavage sites between the two viruses detected in this study, 15 other luchacoviruses, 2 alphacoronaviruses and 3 betacoronaviruses. The genetic comparison of these four regions revealed that two of these regions are located in the C-terminal domain (CTD) of S1 (Figure 3). Region 1 includes the furin cleavage site of SADS-CoV (R-Y-V-R | I, Figure 3A), whose sequence differs in only one residue (residue 458 in Figure 3A) with the respective sequence from luchacoviruses from groups 1 and 2, the divergent variants UMN2020/mouse/2018/USA and P83/plateau pika/China (**<u>S</u>**-Y-V-R | I), and bat CoV HKU2 (**<u>K</u>**-Y-V-R | I). For these variants, a single nucleotide change is sufficient to revert the residue change detected between these viruses (Figure 3B). Compared to group 3 luchacoviruses and the two viruses obtained in this study, it differs in two residues (458 and 459, Figure 3A). In the tertiary structure of SADS-CoV region 1 is located on a beta sheet within the CTD (Figure 3C). Region 2 is highly variable and includes a weakly predicted furin cleavage motif detected in Group 3 luchacoviruses (R-R-A-R | A, ProP score 0.28) and motif F-R | S in W317/red fox/2022/USA consistent with a cathepsin L cleavage site (Figure 3A). In this region divergent luchacoviruses UMN2020/mouse/2018/USA and W291/Bobcat/2021 have highly different sequences (Figure 3A). Region 3 includes the S1/S2 cleavage site, which is observed in betacoronaviruses like SARS-CoV-2 and two rodent betacoronaviruses (ProP scores >0.5), but not in luchacoviruses (Figure 3A). Region 4 includes the S2’cleavage site in which furin cleavage motifs were not predicted for any Luchacovirus. Regions 2, 3 and 4 are located in exposed loops in the tertiary structure of SADS-CoV (Figure 3C).

**Figure 3.**
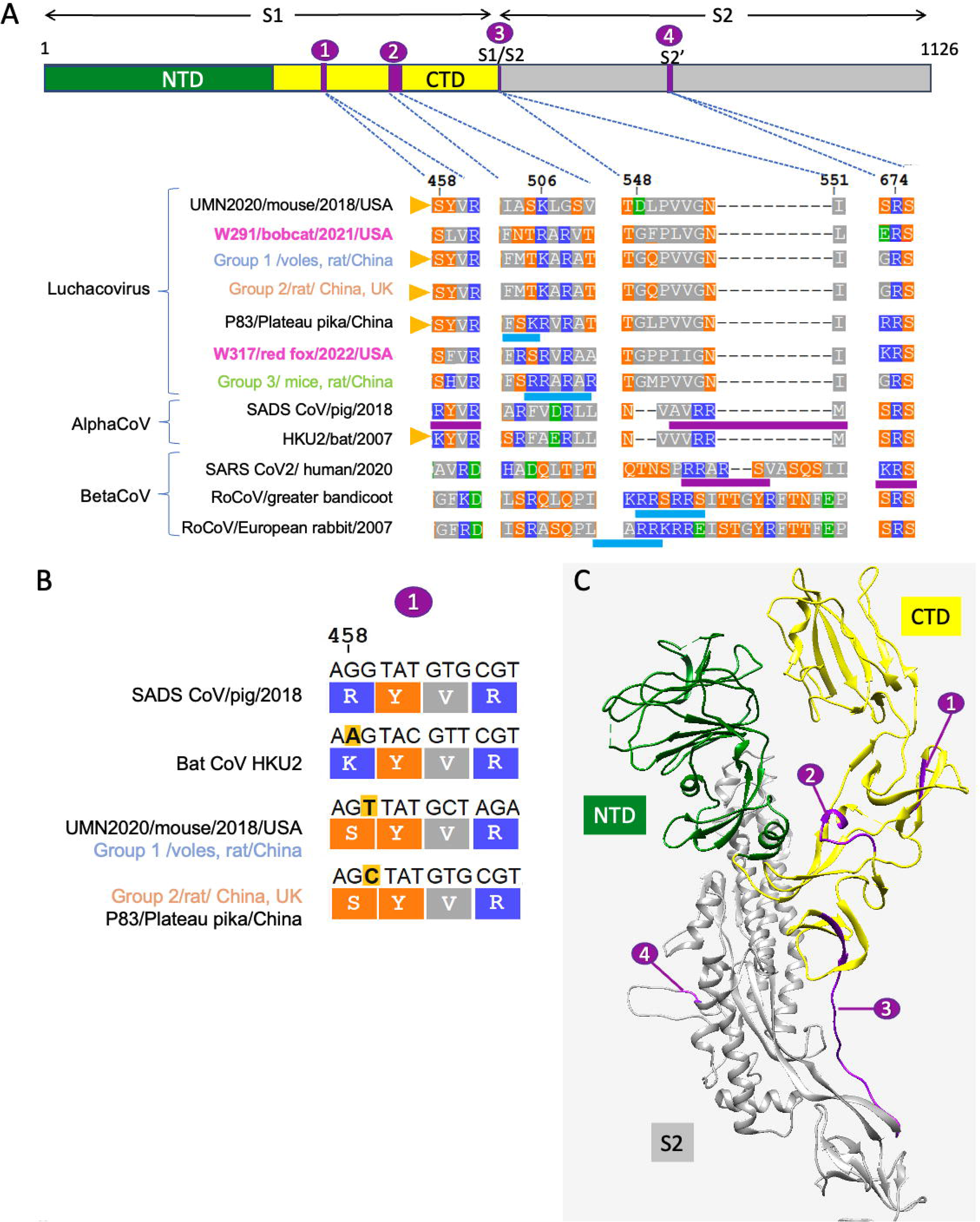
A. Graphical representation of the S protein showing the S1 and S2 domains (top) and multiple sequence alignments (bottom) of four regions of interest within S (shown in violet circles) in which cleavage motifs are found (underlined in blue are the predicted cleavage sites and in violet are experimentally proven furin cleavage sites). The S1 domain is divided into the N-terminal domain (NTD, in green) and the C-terminal domain (CTD, in yellow). In the amino acid alignments polar residues are in orange, acidic residues are in green, basic residues are in blue and nonpolar residues are in grey. In the left side of the alignment the virus name/host/year of collection/place of collection is shown. Above the alignments are the amino acid location of each region based on the S protein sequence of rodent Luchacovirus JC34 (KX964649). From left to right there are four regions of interest: (1) Alphacoronavirus SADS-CoV has a furin cleavage site 94 amino acids upstream of its S1/S2 cleavage site that differs in only one residue when compared to four luchacoviruses and bat CoV HKU2 (shown with yellow triangles), (2) luchacoviruses have a region 37 amino acids upstream of the end of domain S1 that is highly variable and in which there are two different cleavage motifs, (3) luchacoviruses do not have an S1/S2 cleavage region in comparison to SADS-CoV and selected betacoronaviruses (SARS-CoV2 and two rodent coronaviruses) and (4) only SARS-CoV2 has an S2’ cleavage motif. B. Amino acid sequences and the respective codons (on top) of region 1, in which SADS-CoV has a cleavage site and bat CoV HKU2 and the luchacoviruses shown differ by a single residue (458) to that of SADS-CoV (R458S, R458K, respectively), which abrogates the cleavage site. Inspection of the codon sequences shows that only a single nucleotide change (highlighted in yellow) is necessary to revert this residue change. C. Tertiary structure of one of the monomers of the S protein of SADS-CoV showing the four regions in which we found cleavage sites in luchacoviruses or related alpha and betacoronaviruses. For this figure we used the same colors as in A.

The 3’-end of the genome of luchacoviruses has a general organization that includes four structural proteins S, envelope (E), matrix (M), and nucleocapsid (N) and three to five non-structural ORFs, one or two between ORF1b and the S protein (ORF2 and 2b), ORF3 between S and E, ORF6 M and N, ORF8 within N and ORF9 after N (Figure 4). In this region the three luchacovirus groups differed in the number of ORF2, group 1 has ORF2 and 2b, group 2 only has ORF2 and group 3 does not have any ORF2 (Figure 4). The obtained partial genome of W317/red fox/2022 had a main similar organization to group 2 luchacoviruses with only one ORF2 (Figure 4), but differed in that it does not have an ORF8 or its internal transcription signal and ORF9 was shorter than those from the other luchacoviruses, due to a 69-nucleotide deletion (Figure 4). W317/red fox/2022 shared low overall genetic similarity at the 3’-end of the genome when compared to other luchacoviruses (<86%, Figure 5A). In comparison, the luchacoviruses within groups 1, 2, 3 shared an overall high nucleotide similarity (pairwise percentage similarity >94.2%). Divergent luchacoviruses UMN2020/mouse/2018/USA and P83/plateau pika/China had the same genomic organization as luchacoviruses from groups 3 and 2, respectively. However, genetic comparison revealed an overall low similarity throughout the 3’-end of the genome between these luchacoviruses and those in groups 3 and 2 (UMN2020/mouse/2018/USA vs group 3 <82% and P83/plateau pika/China vs group 2 <88%, Figure S2). We did not find any recombination breakpoints in this region of the genome between the 16 luchacoviruses included in the analysis (see Methods). The partial S gene of W291/Bobcat/2021was overall genetically different from other luchacoviruses, including W317/red fox/2022 (<80%, Figure 5B).

**Figure 4.**
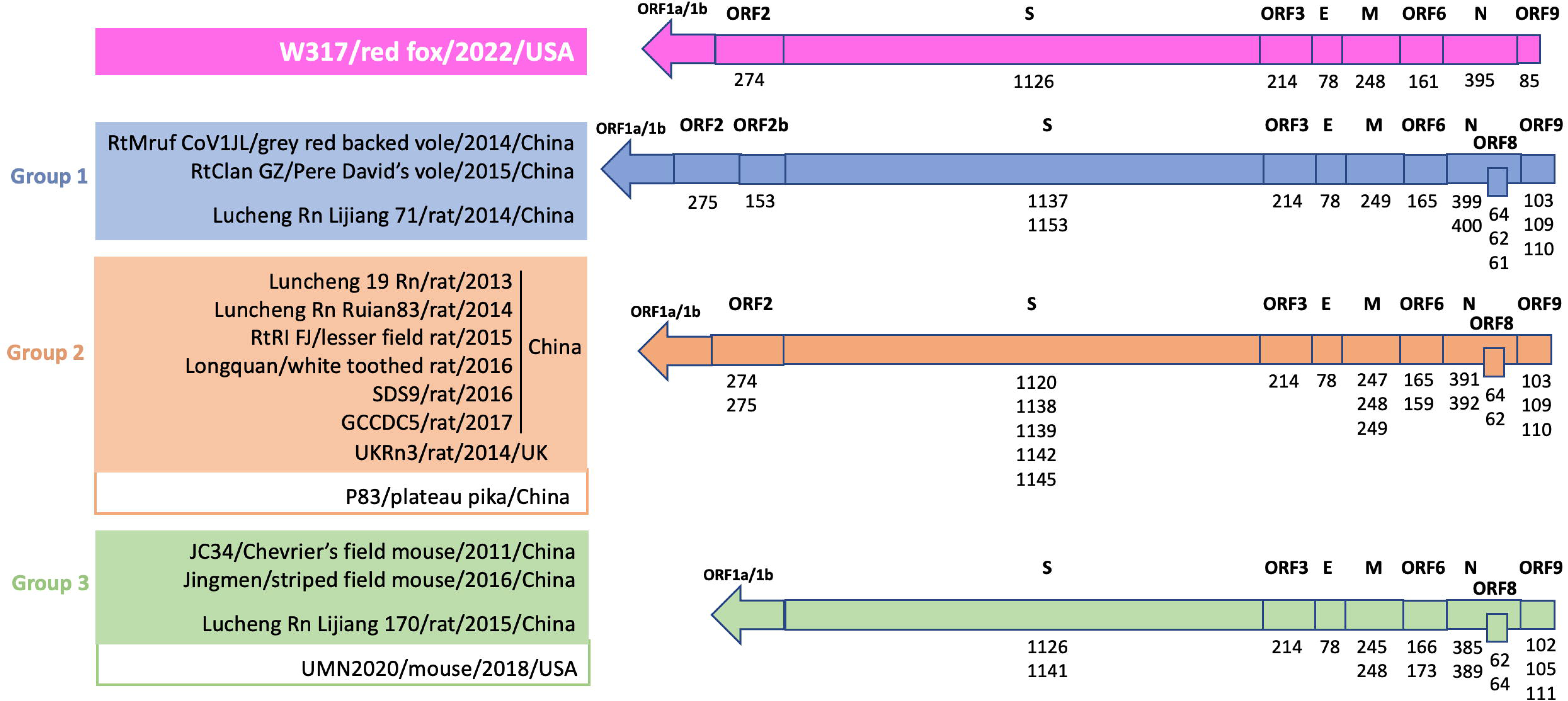
Graphical representation of the 3’-end of the genome of Luchacovirus, that includes the structural (S, E, M, N) and non-structural (ORFs 2, 2b, 6, 8, 9) genes. Sixteen sequences, including the one obtained in this study (on top in fuchsia) are shown. Above each genome representation is the name of each gene, below is the size of the corresponding proteins (number of amino acids). If some proteins varied in size between the viruses, all size values are shown from shortest to longest. On the left side of the figure the name of the virus/host species/year of collection/country of collection of all sequences included are shown. There are three genomic organizations that match the three groups (groups 1, 2 and 3 in blue, orange, and green, respectively) identified in Figure 2. The three genomic organizations differ in the existence of ORFs between ORF1b and S (ORF2 and 2b). The sequence obtained in this study, W317/red fox/2022 (top), has a similar organization to group 2 luchacoviruses as it only has one ORF2 but it differs from other luchacoviruses because it lacks ORF8 and ORF9 is shorter. Divergent luchacoviruses P83/plateau pika/China and UMN2020/mouse/2018/USA have the same genomic organization of viruses from groups 2 and 3, respectively, but do not belong to these groups and are overall genetically different (Figure S2).

**Figure 5.**
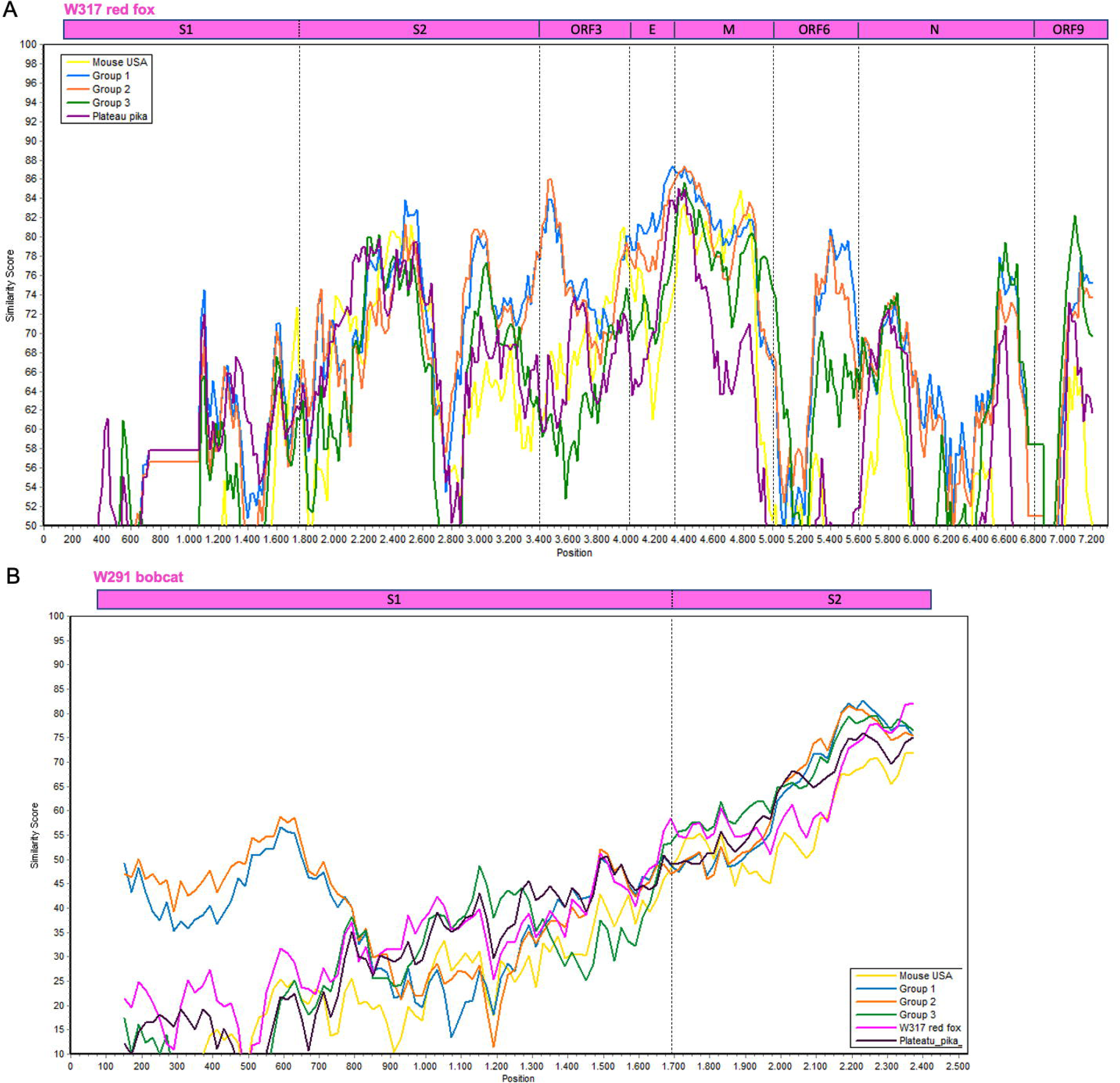
Similarity plot of the nucleotide sequence of (A) the 3’-end of the genome of W317/red fox/2022 (top, used as reference) and (B) a partial region of the S gene of W291/bob cat/2021 (top, used as reference) compared to 15 other luchacoviruses including those in group 1 (in blue), 2 (in orange), and 3 (in green), and divergent luchacoviruses UMN2020/mouse/2018/USA (in yellow) and P83/plateau pika/China (in violet). The location of each gene or gene domain is shown above each the graph. The graphs were constructed using a window of 200bp, a step of 20nt and the kimura 2 parameter distance model.

## Discussion

Coronaviruses are important pathogens that have a wide host range and can result in subclinical infection or cause a variety of diseases in both domesticated and wild mammals. Within the *Alphacoronavirus* genus, the *Luchacovirus* sub-genus includes CoVs reported to date only in rodents and rabbits. In this study we report for the first time luchacoviruses shed in feces and saliva of three species of wild meso-carnivores including bobcat (W291/bobcat/2021), fisher (W145/fisher/2022), and red fox (W317/red fox/2022) and a feral domestic cat (DC6837 F and S/domestic cat/2022) living freely in NYC (Figure 1). The phylogeny of the S1 domain of the S protein and the general genomic organization of the rodent and rabbit luchacoviruses reported to date showed that there are three main groups (Groups 1-3, Figures 2 and 4, Figure S1), two of which were already observed using fewer sequences (57). Phylogenetic and genetic comparison of the S1 domain of two of the viruses obtained in this study (W291/bobcat/2021 and W317/red fox/2022) revealed that this region is genetically different from the luchacoviruses reported to date in rodents and rabbits (<75%, Figure 5) and that these viruses did not group within the three groups that comprise most of the luchacoviruses reported to date (Figure 2). Although in general CoVs have highly variable S1 regions, further genetic comparison of the 3’-end of the genome revealed that luchacoviruses within the three established groups are highly similar (>94.2% similarity) but W317/red fox/2022 is overall divergent (<86% nucleotide similarity, Figure 5). Likewise, the genetic organization of the 3’-end of the genome of W317/red fox/2022 had certain differences with the other luchacoviruses including a shorter ORF9 and the absence of ORF8 (Figure 4). We found that the only available luchacovirus reported from a mouse in the U.S. (UMN2020/mouse/2018/USA) has the same genetic organization as Group 3 luchacoviruses, which includes viruses detected in rodents in China (Figure 4). These results support the idea that, with the sequences available to date, at least two of the viruses reported in this study (W291/bobcat/2021 and W317/red fox/2022), from which we obtained partial genome sequences, are genetically different from other luchacoviruses reported in rodents and rabbits.

We only detected luchacovirus RNA in feces and saliva. Therefore, it is possible that these may be viruses from the prey consumed by these animals, as has been shown for insectivorous bats which shed various insect viruses in their feces (58). However, one of the luchacoviruses was detected in feces and saliva of a feral cat living in NYC (Figure 1). A previous study of cat colonies in NYC found that 94.3% of sampled cat scat contained cow, chicken, and fish taxa, indicating that most of the diet of these cats consists of cat food provided by humans (59) Additionally, it has been reported that enteric viruses like norovirus, astrovirus, and rotavirus can replicate in salivary glands and be transmitted through saliva (60). The sequences obtained from this feral cat (DC6837 F and S/domestic cat/2022) were identical to the one obtained from the red fox (W317/red fox/2022, Figure 1), from which we were able to sequence the complete 3’-end of the genome (Figure 4). Our phylogenetic analyses of the three structural genes (E, M, and N) revealed that W317/red fox/2022 did not cluster with any of the three luchacovirus groups, and only grouped with the luchacovirus UMN2020/mouse/2018/USA, which was detected in a mouse bought at a pet store in the U.S. (61) in the M gene tree (Figure S1). However, a genetic comparison of the M gene between these two viruses showed that their percentage nucleotide similarity was low (<86%, Figure 5). Overall, the luchacoviruses reported from the US, including those reported in this study (W291/bobcat/2021 and W317/red fox/2022) and the one previously reported from a mouse (UMN2020/mouse/2018/USA) were genetically highly variable and did not group within any luchacovirus group (Figure 2, Figure S1). Nevertheless, as stray cats also feed occasionally from mice (59), and cat food stations in urban areas attract rodents (especially house mice (62), it is necessary to screen both rodents and feral cats to unveil the importance of these interspecies gatherings as well as predator-prey interactions in the epidemiology and possible cross-species transmission of luchacoviruses. Moreover, since in the phylogeny of the partial RdRp gene W317/red fox/2022, W145/fisher/2022, and DC6837 F and S /domestic cat/2022 grouped with viruses of vole and rabbit from which only partial sequences exist (Figure 1), additional sequences of luchacoviruses from these species will also help clarify the host range of these viruses. Whether luchacovirus can infect meso-carnivores, or the detected viral RNA was derived from their prey, our results reveal that luchacoviruses circulating in the U.S. are more genetically diverse than those reported in Asia and Europe to date.

Some CoVs have wide host ranges, possibly due to the use of protein receptors like angiotensin converting enzyme 2 (ACE-2), dipeptidyl peptidase 4 (DPP4) or aminopeptidase N (APN), that are highly conserved between diverse species (44). As the S1 domain of the S protein of CoVs interacts with the receptor, signatures of viral adaptation to the receptor of different species have been identified in this region which has helped increase our understanding of the molecular mechanisms of the adaptation and emergence of CoVs in various species (12). Much less is known about the relation between the cleavage of S and the emergence of pathogenic CoV variants, however the presence of a unique cleavage site in the S1/S2 of SARS-CoV2, that to date has not been found in closely related viruses (63), has highlighted the importance of studying the mechanisms CoVs may use to acquire functional cleavage (22). For example, genetic comparison of the S1/S2 region between SARS-CoV-2 and closely related CoVs from European bats revealed that a single nucleotide substitution is sufficient to produce a functional furin cleavage site identical to that of SARS-CoV-2 in two bat CoV variants, an essential molecular determinant of their zoonotic potential (21). However, this substitution alone is not enough to enable furin cleavage in similar bat CoVs, possibly because it is in a short loop that may not be accessible to the protease (64). The Luchacovirus clade is closely related to SADS-CoV and bat CoV HKU2 (Figure 1 and 2), which are also characterized for having a recombinant S gene that is more similar to betacoronaviruses than alphacoronaviruses (65). To date the receptor of luchacoviruses, SADS-CoV or bat CoV HKU2 remains unknown (66); in this study we directed our genetic analysis of the S protein on finding regions with putative cleavage sites and the genetic comparison of these regions with other CoVs like SADS-CoV (11) and SARS-CoV-2 in which the cleavage of S has been experimentally studied.

SADS-CoV is a highly pathogenic CoV that infects pigs and has a unique cleavage site (R-Y-V-R | I) in the S1 domain (region 1 in Figure 3) that is essential for cell fusion and may have an impact in the pathogenicity of this virus and its observed broad cell range (67). Comparative genetic analyses have shown that bat CoV HKU2 is closely related to SADS-CoV, suggesting that bat CoVs may also be involved in the emergence of pathogenic CoVs of veterinary importance (55). It has been previously shown that a substitution of a single residue in this cleavage site (residue 458 in Figure 3) is enough to abrogate the cleavage of S (11). In this study we show that several luchacoviruses, including those in groups 1 and 2 and divergent variants UMN2020/mouse/2018/USA and P83/plateau pika/China, have a cleavage motif in this region that differ only in this essential residue to that of SADS-CoV (R458S, Figure 3A) and that a single nucleotide substitution is sufficient to revert this residue (Figure 3B). Likewise, bat CoV HKU2 also differs only in this residue when compared to SADS-CoV (R458K, Figures 3A and B). However, we show that unlike SADS-CoV and bat CoV HKU2, luchacoviruses do not have an S1/S2 cleavage site (Figure 3A). As cleavage in both S1 and S1/S2 sites is required for cleavage and cell fusion of SADS-CoV (11), more than a single non-synonymous change would be necessary for luchacoviruses to acquire a cleavage mechanism like the one of pathogenic SADS-CoV. In contrast to luchacoviruses, most rodent betacoronaviruses have a predicted S1/S2 cleavage site (78% of available S sequences, (22)), more than 13 times of what was predicted for bat betacoronaviruses (6% of available sequences, (22)). *In vitro* experiments have shown that SADS-CoV can successfully infect and replicate in several rodent cell lines (66); therefore, surveillance of rodents, meso-carnivores, pigs, and bats targeting luchacoviruses, rodent betacoronaviruses and SADS-CoV and characterizing their S genes is essential to detect if there is any recombination or variation in these key regions and its relationship with pathogenicity and host range.

A previous study found that Luchacovirus JC34 had a predicted furin cleavage site (R-R-A-R | A) in the S1 domain similar to that of the S1/S2 cleavage site SARS-CoV2 (22). In this study we identified two additional luchacoviruses (all from group 3, Figure 2) that also have this cleavage site (Figure 3). As it has been reported that this cleavage motif is not cleaved by human furin (27), we conducted a genetic comparison of this region among luchacoviruses, which revealed that this region is overall genetically highly variable. Aside from the furin cleavage motif, we also identified possible motifs for cathepsin L or V cleavage (F-R | S) in divergent luchacovirus P83/plateau pika/China (Figure 3A). As cleavage motifs need to be accessible to proteases, we mapped this region in the tertiary structure of SADS-CoV and show that it is located in an exposed loop (region 2 in Figure 3), and it is, like region 3 (S1/S2 cleavage site) and region 4 (S2’ cleavage site, Figure 3C). Therefore, identifying cleavage sites in divergent CoVs together with their mapping on crystalized tertiary structures can help find regions that may be critical in the context of the virus-host interactions, and which can be targeted for future laboratory testing. Aside from the two viruses reported in this study, two additional luchacoviruses (UMN2020/mouse/2018/USA (61) and P83/plateau pika/China (68)) are highly divergent (Figure 2, Figure S2), thus showing that luchacoviruses are highly diverse and surveillance targeting both conserved and variable regions like the S1 may be essential to understand CoV diversity and evolution. With the results shown in this study we increased the dataset of sequences from luchacoviruses extending the known host range of these viruses and showing possible regions of interest in the S protein for future surveillance.

We did not find recombination breakpoints in the genomic region analyzed between the luchacovirus sequences available to date. However possible recombination breakpoints have been detected in the ORF1ab between viruses of groups 1, 2 and 3 (57). This may indicate that these viruses are infecting different hosts at the same time. In this study we are reporting divergent luchacoviruses in three species of widely spread meso-carnivore, whose diet include rabbits and rodents, and thus it would be interesting to assess if the predator-prey interactions play a role in the spread of CoVs in wild populations. However, we did not find luchacoviruses in the rodent and rabbit samples screened in this study (Table S1) possibly due to our smaller sample size (36 samples) compared to that of the meso-carnivore (300 samples) in which we found luchacoviruses in 1% (3 positive samples). In previous studies from China (26) and Europe (69) it has been shown that luchacoviruses are less prevalent than rodent and rabbit betacoronaviruses (0.8% to 1.1% of positive samples were luchacoviruses compared to 6.1 to 11.8% for rodent betacoronaviruses). It is necessary to continue the surveillance of wildlife (including meso-carnivores, rodents, and lagomorphs) in the U.S. to increase our knowledge of the viral diversity hosted in native species. Since all the recovered viral sequences herein reported were obtained from fecal samples, we show it is possible to perform a successful surveillance of alphacoronaviruses in meso-carnivore in a non-invasive manner.

## Acknowledgements

This work was funded by the Cornell Public and Ecosystem Health Impact Award and Alfred P. Sloan Foundation grant number G-2022-19422. Capacity for coronavirus discovery screening was supported (FOA PAR-18-604) by the US Food and Drug Administration’s Veterinary Laboratory Investigation and Response Network under grant 1U18FD006993-01.

The authors thank Melisa A. Fadden for assistance with sample storage and retrieval; Christian Lange for assistance with screening protocols; Guillaume Reboul and Joe Flint for technical support. RNA-seq was conducted by scientists in the Transcriptional Regulation and Expression Facility at Cornell University (RRID: SCR_022532).

## Data Availability

Sequences obtained in this study are deposited in GenBank (accession numbers OQ756331-35 and OR428266-7).

## References

1. Siddell SG, Walker PJ, Lefkowitz EJ, Mushegian AR, Adams MJ, Dutilh BE, Gorbalenya AE, Harrach B, Harrison RL, Junglen S, Knowles NJ, Kropinski AM, Krupovic M, Kuhn JH, Nibert M, Rubino L, Sabanadzovic S, Sanfaçon H, Simmonds P, Varsani A, Zerbini FM, Davison AJ. 2019. Additional changes to taxonomy ratified in a special vote by the International Committee on Taxonomy of Viruses (October 2018). Arch Virol 164:943–946.

2. Cui J, Li F, Shi Z-L. 2019. Origin and evolution of pathogenic coronaviruses. Nature Reviews Microbiology 17:181–192.

3. Li F. 2015. Receptor recognition mechanisms of coronaviruses: a decade of structural studies. J Virol 89:1954–64.

4. Belouzard S, Millet JK, Licitra BN, Whittaker GR. 2012. Mechanisms of coronavirus cell entry mediated by the viral spike protein. Viruses 4:1011–33.

5. Millet JK, Whittaker GR. 2015. Host cell proteases: Critical determinants of coronavirus tropism and pathogenesis. Virus Res 202:120–34.

6. Belouzard S, Chu VC, Whittaker GR. 2009. Activation of the SARS coronavirus spike protein via sequential proteolytic cleavage at two distinct sites. Proc Natl Acad Sci U S A 106:5871–6.

7. Wu Y, Zhao S. 2020. Furin cleavage sites naturally occur in coronaviruses. Stem Cell Res 50:102115.

8. Chan YA, Zhan SH. 2022. The Emergence of the Spike Furin Cleavage Site in SARS-CoV-2. Mol Biol Evol 39.

9. Jaimes JA, Whittaker GR. 2018. Feline coronavirus: Insights into viral pathogenesis based on the spike protein structure and function. Virology 517:108–121.

10. Gong L, Li J, Zhou Q, Xu Z, Chen L, Zhang Y, Xue C, Wen Z, Cao Y. 2017. A New Bat-HKU2-like Coronavirus in Swine, China, 2017. Emerg Infect Dis 23:1607–9.

11. Kim J, Yoon J, Park JE. 2022. Furin cleavage is required for swine acute diarrhea syndrome coronavirus spike protein-mediated cell - cell fusion. Emerg Microbes Infect 11:2176–2183.

12. Li F. 2016. Structure, Function, and Evolution of Coronavirus Spike Proteins. Annual Review of Virology 3:237–261.

13. Jaimes JA, Millet JK, Stout AE, Andre NM, Whittaker GR. 2020. A Tale of Two Viruses: The Distinct Spike Glycoproteins of Feline Coronaviruses. Viruses 12.

14. Choe Y, Leonetti F, Greenbaum DC, Lecaille F, Bogyo M, Brömme D, Ellman JA, Craik CS. 2006. Substrate profiling of cysteine proteases using a combinatorial peptide library identifies functionally unique specificities. J Biol Chem 281:12824–32.

15. Yang TJ, Chang YC, Ko TP, Draczkowski P, Chien YC, Chang YC, Wu KP, Khoo KH, Chang HW, Hsu SD. 2020. Cryo-EM analysis of a feline coronavirus spike protein reveals a unique structure and camouflaging glycans. Proc Natl Acad Sci U S A 117:1438–1446.

16. Tang T, Jaimes JA, Bidon MK, Straus MR, Daniel S, Whittaker GR. 2021. Proteolytic Activation of SARS-CoV-2 Spike at the S1/S2 Boundary: Potential Role of Proteases beyond Furin. ACS Infect Dis 7:264–272.

17. Hatmal MM, Alshaer W, Al-Hatamleh MAI, Hatmal M, Smadi O, Taha MO, Oweida AJ, Boer JC, Mohamud R, Plebanski M. 2020. Comprehensive Structural and Molecular Comparison of Spike Proteins of SARS-CoV-2, SARS-CoV and MERS-CoV, and Their Interactions with ACE2. Cells 9.

18. Licitra BN, Millet JK, Regan AD, Hamilton BS, Rinaldi VD, Duhamel GE, Whittaker GR. 2013. Mutation in spike protein cleavage site and pathogenesis of feline coronavirus. Emerg Infect Dis 19:1066–73.

19. Johnson BA, Xie X, Bailey AL, Kalveram B, Lokugamage KG, Muruato A, Zou J, Zhang X, Juelich T, Smith JK, Zhang L, Bopp N, Schindewolf C, Vu M, Vanderheiden A, Winkler ES, Swetnam D, Plante JA, Aguilar P, Plante KS, Popov V, Lee B, Weaver SC, Suthar MS, Routh AL, Ren P, Ku Z, An Z, Debbink K, Diamond MS, Shi PY, Freiberg AN, Menachery VD. 2021. Loss of furin cleavage site attenuates SARS-CoV-2 pathogenesis. Nature 591:293–299.

20. Millet JK, Whittaker GR. 2014. Host cell entry of Middle East respiratory syndrome coronavirus after two-step, furin-mediated activation of the spike protein. Proc Natl Acad Sci U S A 111:15214–9.

21. Sander AL, Moreira-Soto A, Yordanov S, Toplak I, Balboni A, Ameneiros RS, Corman V, Drosten C, Drexler JF. 2022. Genomic determinants of Furin cleavage in diverse European SARS-related bat coronaviruses. Commun Biol 5:491.

22. Stout AE, Millet JK, Stanhope MJ, Whittaker GR. 2021. Furin cleavage sites in the spike proteins of bat and rodent coronaviruses: Implications for virus evolution and zoonotic transfer from rodent species. One Health 13:100282.

23. Tsoleridis T, Chappell JG, Onianwa O, Marston DA, Fooks AR, Monchatre-Leroy E, Umhang G, Müller MA, Drexler JF, Drosten C, Tarlinton RE, McClure CP, Holmes EC, Ball JK. 2019. Shared Common Ancestry of Rodent Alphacoronaviruses Sampled Globally. Viruses 11.

24. Wu Z, Lu L, Du J, Yang L, Ren X, Liu B, Jiang J, Yang J, Dong J, Sun L, Zhu Y, Li Y, Zheng D, Zhang C, Su H, Zheng Y, Zhou H, Zhu G, Li H, Chmura A, Yang F, Daszak P, Wang J, Liu Q, Jin Q. 2018. Comparative analysis of rodent and small mammal viromes to better understand the wildlife origin of emerging infectious diseases. Microbiome 6:178.

25. Tsoleridis T, Onianwa O, Horncastle E, Dayman E, Zhu M, Danjittrong T, Wachtl M, Behnke JM, Chapman S, Strong V, Dobbs P, Ball JK, Tarlinton RE, McClure CP. 2016. Discovery of Novel Alphacoronaviruses in European Rodents and Shrews. Viruses 8:84.

26. Ge XY, Yang WH, Zhou JH, Li B, Zhang W, Shi ZL, Zhang YZ. 2017. Detection of alpha- and betacoronaviruses in rodents from Yunnan, China. Virol J 14:98.

27. Choi A, Singleton DT, Stout AE, Millet JK, Whittaker GR. 2021. In vitro and computational analysis of the putative furin cleavage site (RRARS) in the divergent spike protein of the rodent coronavirus AcCoV-JC34 (sub-genus luchacovirus). bioRxiv doi:10.1101/2021.12.16.473025:2021.12.16.473025.

28. Ray JC. 2000. Mesocarnivores of Northeastern North America: Status and Conservation Issues. WCS Working Papers No 15. Wildlife Conservation Society, Bronx, New York.

29. Prange S, Gehrt SD, Wiggers EP. 2003. Demographic Factors Contributing to High Raccoon Densities in Urban Landscapes. The Journal of Wildlife Management 67:324–333.

30. Sabeta CT, Mansfield KL, McElhinney LM, Fooks AR, Nel LH. 2007. Molecular epidemiology of rabies in bat-eared foxes (Otocyon megalotis) in South Africa. Virus Res 129:1–10.

31. Jessup DA. 2004. The welfare of feral cats and wildlife. J Am Vet Med Assoc 225:1377–83.

32. Grieco V, Crepaldi P, Giudice C, Roccabianca P, Sironi G, Brambilla E, Magistrelli S, Ravasio G, Granatiero F, Invernizzi A, Caniatti M. 2021. Causes of Death in Stray Cat Colonies of Milan: A Five-Year Report. Animals 11:3308.

33. Attipa C, Gunn-Moore D, Mazeri S, Epaminondas D, Lyraki M, Hardas A, Loukaidou S, Gentil M. 2023. Concerning feline infectious peritonitis outbreak in Cyprus. Veterinary Record 192:449–450.

34. Roemer GW, Gompper ME, Van Valkenburgh B. 2009. The Ecological Role of the Mammalian Mesocarnivore. BioScience 59:165–173.

35. Vlasova AN, Halpin R, Wang S, Ghedin E, Spiro DJ, Saif LJ. 2011. Molecular characterization of a new species in the genus Alphacoronavirus associated with mink epizootic catarrhal gastroenteritis. J Gen Virol 92:1369–1379.

36. Murray J, Kiupel M, Maes RK. 2010. Ferret coronavirus-associated diseases. Vet Clin North Am Exot Anim Pract 13:543–60.

37. Tekes G, Thiel HJ. 2016. Feline Coronaviruses: Pathogenesis of Feline Infectious Peritonitis. Adv Virus Res 96:193–218.

38. Erles K, Brownlie J. 2008. Canine respiratory coronavirus: an emerging pathogen in the canine infectious respiratory disease complex. Vet Clin North Am Small Anim Pract 38:815–25, viii.

39. Dong BQ, Liu W, Fan XH, Vijaykrishna D, Tang XC, Gao F, Li LF, Li GJ, Zhang JX, Yang LQ, Poon LL, Zhang SY, Peiris JS, Smith GJ, Chen H, Guan Y. 2007. Detection of a novel and highly divergent coronavirus from asian leopard cats and Chinese ferret badgers in Southern China. J Virol 81:6920–6.

40. Kim YI, Kim SG, Kim SM, Kim EH, Park SJ, Yu KM, Chang JH, Kim EJ, Lee S, Casel MAB, Um J, Song MS, Jeong HW, Lai VD, Kim Y, Chin BS, Park JS, Chung KH, Foo SS, Poo H, Mo IP, Lee OJ, Webby RJ, Jung JU, Choi YK. 2020. Infection and Rapid Transmission of SARS-CoV-2 in Ferrets. Cell Host Microbe 27:704–709.e2.

41. Oude Munnink BB, Sikkema RS, Nieuwenhuijse DF, Molenaar RJ, Munger E, Molenkamp R, van der Spek A, Tolsma P, Rietveld A, Brouwer M, Bouwmeester-Vincken N, Harders F, Hakze-van der Honing R, Wegdam-Blans MCA, Bouwstra RJ, GeurtsvanKessel C, van der Eijk AA, Velkers FC, Smit LAM, Stegeman A, van der Poel WHM, Koopmans MPG. 2021. Transmission of SARS-CoV-2 on mink farms between humans and mink and back to humans. Science 371:172–177.

42. Vlasova AN, Diaz A, Damtie D, Xiu L, Toh T-H, Lee JS-Y, Saif LJ, Gray GC. 2021. Novel Canine Coronavirus Isolated from a Hospitalized Patient With Pneumonia in East Malaysia. Clinical Infectious Diseases 74:446–454.

43. Zehr JD, Pond SLK, Martin DP, Ceres K, Whittaker GR, Millet JK, Goodman LB, Stanhope MJ. 2022. Recent Zoonotic Spillover and Tropism Shift of a Canine Coronavirus Is Associated with Relaxed Selection and Putative Loss of Function in NTD Subdomain of Spike Protein. Viruses 14:853.

44. Olarte-Castillo XA, Dos Remedios JF, Heeger F, Hofer H, Karl S, Greenwood AD, East ML. 2021. The virus-host interface: Molecular interactions of Alphacoronavirus-1 variants from wild and domestic hosts with mammalian aminopeptidase N. Mol Ecol 30:2607–2625.

45. Campbell SJ, Ashley W, Gil-Fernandez M, Newsome TM, Di Giallonardo F, Ortiz-Baez AS, Mahar JE, Towerton AL, Gillings M, Holmes EC, Carthey AJR, Geoghegan JL. 2020. Red fox viromes in urban and rural landscapes. Virus Evol 6:veaa065.

46. Kalantar KL, Carvalho T, de Bourcy CFA, Dimitrov B, Dingle G, Egger R, Han J, Holmes OB, Juan YF, King R, Kislyuk A, Lin MF, Mariano M, Morse T, Reynoso LV, Cruz DR, Sheu J, Tang J, Wang J, Zhang MA, Zhong E, Ahyong V, Lay S, Chea S, Bohl JA, Manning JE, Tato CM, DeRisi JL. 2020. IDseq-An open source cloud-based pipeline and analysis service for metagenomic pathogen detection and monitoring. Gigascience 9.

47. Sievers F, Wilm A, Dineen D, Gibson TJ, Karplus K, Li W, Lopez R, McWilliam H, Remmert M, Söding J, Thompson JD, Higgins DG. 2011. Fast, scalable generation of high-quality protein multiple sequence alignments using Clustal Omega. Mol Syst Biol 7:539.

48. Tamura K, Stecher G, Kumar S. 2021. MEGA11: Molecular Evolutionary Genetics Analysis Version 11. Mol Biol Evol 38:3022–3027.

49. Edgar RC. 2004. MUSCLE: multiple sequence alignment with high accuracy and high throughput. Nucleic Acids Res 32:1792–7.

50. Lole KS, Bollinger RC, Paranjape RS, Gadkari D, Kulkarni SS, Novak NG, Ingersoll R, Sheppard HW, Ray SC. 1999. Full-length human immunodeficiency virus type 1 genomes from subtype C-infected seroconverters in India, with evidence of intersubtype recombination. J Virol 73:152–60.

51. Martin DP, Varsani A, Roumagnac P, Botha G, Maslamoney S, Schwab T, Kelz Z, Kumar V, Murrell B. 2021. RDP5: a computer program for analyzing recombination in, and removing signals of recombination from, nucleotide sequence datasets. Virus Evol 7:veaa087.

52. Martin D, Rybicki E. 2000. RDP: detection of recombination amongst aligned sequences. Bioinformatics 16:562–3.

53. Peacock TP, Goldhill DH, Zhou J, Baillon L, Frise R, Swann OC, Kugathasan R, Penn R, Brown JC, Sanchez-David RY, Braga L, Williamson MK, Hassard JA, Staller E, Hanley B, Osborn M, Giacca M, Davidson AD, Matthews DA, Barclay WS. 2021. The furin cleavage site in the SARS-CoV-2 spike protein is required for transmission in ferrets. Nature Microbiology 6:899–909.

54. Duckert P, Brunak S, Blom N. 2004. Prediction of proprotein convertase cleavage sites. Protein Eng Des Sel 17:107–12.

55. Yu J, Qiao S, Guo R, Wang X. 2020. Cryo-EM structures of HKU2 and SADS-CoV spike glycoproteins provide insights into coronavirus evolution. Nature Communications 11:3070.

56. Goddard TD, Huang CC, Meng EC, Pettersen EF, Couch GS, Morris JH, Ferrin TE. 2018. UCSF ChimeraX: Meeting modern challenges in visualization and analysis. Protein Sci 27:14–25.

57. Wang W, Lin XD, Zhang HL, Wang MR, Guan XQ, Holmes EC, Zhang YZ. 2020. Extensive genetic diversity and host range of rodent-borne coronaviruses. Virus Evol 6:veaa078.

58. Li L, Victoria JG, Wang C, Jones M, Fellers GM, Kunz TH, Delwart E. 2010. Bat Guano Virome: Predominance of Dietary Viruses from Insects and Plants plus Novel Mammalian Viruses. Journal of Virology 84:6955–6965.

59. Plimpton LD, Henger CS, Munshi-South J, Tufts D, Kross S, Diuk-Wasser M. 2021. Use of molecular scatology to assess the diet of feral cats living in urban colonies. Journal of Urban Ecology 7.

60. Ghosh S, Kumar M, Santiana M, Mishra A, Zhang M, Labayo H, Chibly AM, Nakamura H, Tanaka T, Henderson W, Lewis E, Voss O, Su Y, Belkaid Y, Chiorini JA, Hoffman MP, Altan-Bonnet N. 2022. Enteric viruses replicate in salivary glands and infect through saliva. Nature 607:345–350.

61. Fay EJ, Balla KM, Roach SN, Shepherd FK, Putri DS, Wiggen TD, Goldstein SA, Pierson MJ, Ferris MT, Thefaine CE, Tucker A, Salnikov M, Cortez V, Compton SR, Kotenko SV, Hunter RC, Masopust D, Elde NC, Langlois RA. 2022. Natural rodent model of viral transmission reveals biological features of virus population dynamics. J Exp Med 219.

62. Hawkins CC, Grant WE, Longnecker MT. 1999. Effect of Subsidized house cats on California birds and rodents. 1999 Transactions of the Western Section of the Wildlife Society 35:29–33.

63. Coutard B, Valle C, de Lamballerie X, Canard B, Seidah NG, Decroly E. 2020. The spike glycoprotein of the new coronavirus 2019-nCoV contains a furin-like cleavage site absent in CoV of the same clade. Antiviral Res 176:104742.

64. Tan CCS, Trew J, Peacock TP, Mok KY, Hart C, Lau K, Ni D, Orme CDL, Ransome E, Pearse WD, Coleman CM, Bailey D, Thakur N, Quantrill JL, Sukhova K, Richard D, Kahane L, Woodward G, Bell T, Worledge L, Nunez-Mino J, Barclay W, van Dorp L, Balloux F, Savolainen V. 2023. Genomic screening of 16 UK native bat species through conservationist networks uncovers coronaviruses with zoonotic potential. Nature Communications 14:3322.

65. Fu X, Fang B, Liu Y, Cai M, Jun J, Ma J, Bu D, Wang L, Zhou P, Wang H, Zhang G. 2018. Newly emerged porcine enteric alphacoronavirus in southern China: Identification, origin and evolutionary history analysis. Infect Genet Evol 62:179–187.

66. Edwards CE, Yount BL, Graham RL, Leist SR, Hou YJ, Dinnon KH, 3rd, Sims AC, Swanstrom J, Gully K, Scobey TD, Cooley MR, Currie CG, Randell SH, Baric RS. 2020. Swine acute diarrhea syndrome coronavirus replication in primary human cells reveals potential susceptibility to infection. Proc Natl Acad Sci U S A 117:26915–26925.

67. Guo Y-Y, Wang P-H, Pan Y-Q, Shi R-Z, Li Y-Q, Guo F, Xing L. 2021. The Characteristics of Spike Glycoprotein Gene of Swine Acute Diarrhea Syndrome Coronavirus Strain CH/FJWT/2018 Isolated in China. Frontiers in Veterinary Science 8.

68. Zhu W, Yang J, Lu S, Jin D, Wu S, Pu J, Luo X-l, Liu L, Li Z, Xu J. 2021. Discovery and Evolution of a Divergent Coronavirus in the Plateau Pika From China That Extends the Host Range of Alphacoronaviruses. Frontiers in Microbiology 12.

69. Monchatre-Leroy E, Boué F, Boucher JM, Renault C, Moutou F, Ar Gouilh M, Umhang G. 2017. Identification of Alpha and Beta Coronavirus in Wildlife Species in France: Bats, Rodents, Rabbits, and Hedgehogs. Viruses 9.

