## Supplemental data for "Detection and characterization of novel luchacoviruses, genus *Alphacoronavirus*, shed in saliva and feces of meso-carnivores in the northeastern United States"

**Table S1**. Rabbit and rodent screening results according to sample type and species.

| Species | Collection year | Sample type | Number of samples | Number of positive samples |
| --- | --- | --- | --- | --- |
| Chipmunk | 2021 | Feces | 6 | 0 |
| Chipmunk | 2021 | Lung | 6 | 0 |
| Eastern cottontail | 2021 | Feces | 4 | 0 |
| Eastern cottontail | 2021 | Lung | 4 | 0 |
| Gray squirrel | 2021 | feces | 6 | 0 |
| Gray squirrel | 2021 | Lung | 6 | 0 |
| Porcupine | 2021 | Feces | 2 | 0 |
| Porcupine | 2021 | Lung | 2 | 0 |
|  |  |  | 36 | 0 |

**Figure S1.** Maximum likelihood tree of the complete nucleotide sequence of the E (A), M (B), and N (C) genes of 16 luchacoviruses including one obtained in this study (in fuchsia). The groups observed in the S1 tree (Figure 2) are also present in all trees based on these genes (Group 1 in blue, Group 2 in orange, Group 3 in green). The luchacovirus detected in this study on a red fox in the U.S. does not cluster with any of these groups and only in the tree of the M gene (B) it groups with the sequence of the luchacovirus obtained from a mouse in the U.S. For all sequences included, the virus name, host species, year, place of collection, and Genbank accession number are shown at the tip of the tree. Numbers at the branches indicate bootstrap percentage values from 1000 replicates. Branches with bootstrap support of <50 were collapsed. Nucleotide substitution model used: GTR+I+G for M and N genes and HKY + I for the E gene.

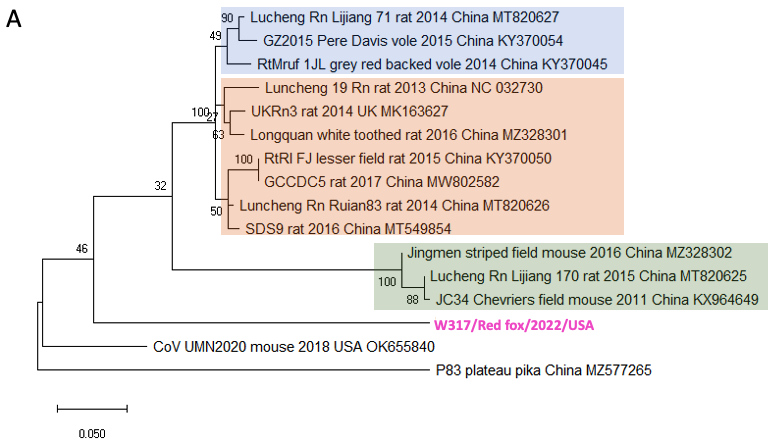

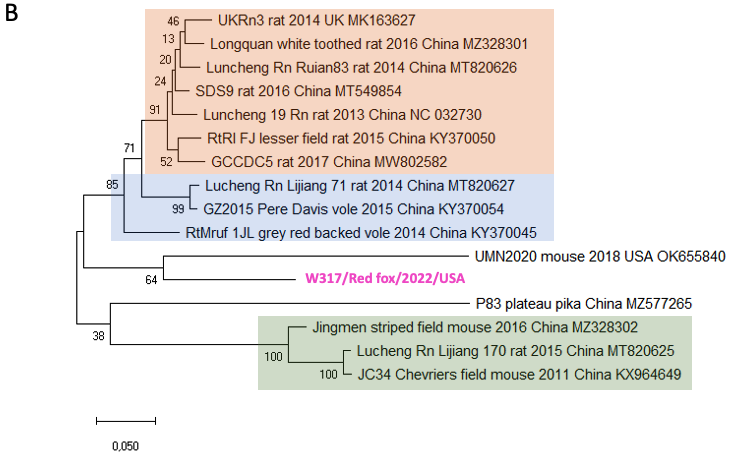

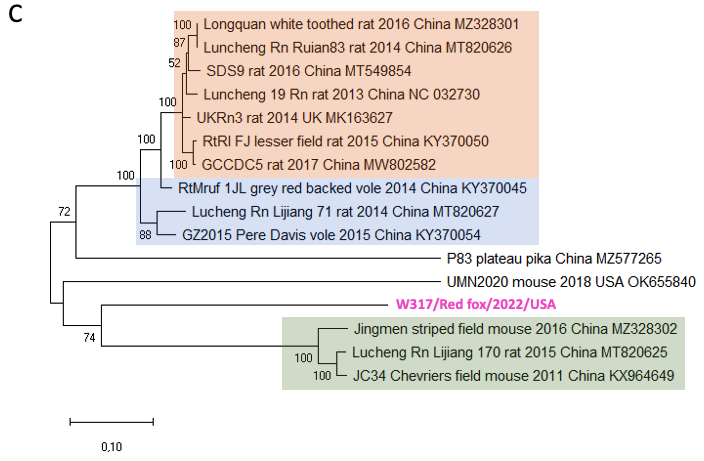

**Figure S2**. Similarity plots comparing the nucleotide sequence of the 3’-end of the genome of UMN2020/mouse/2018/USA (in violet) with group 3 luchacoviruses (in green, top) and P83/plateau pika/China (red) with group 2 luchacoviruses (in orange, bottom). The graphs were constructed using a window of 200bp, a step 20nt and the kimura 2 parameter distance model. and Kimura-2-parameter distance model with a sliding window of 200 nucleotides and a step size of 20 bases.

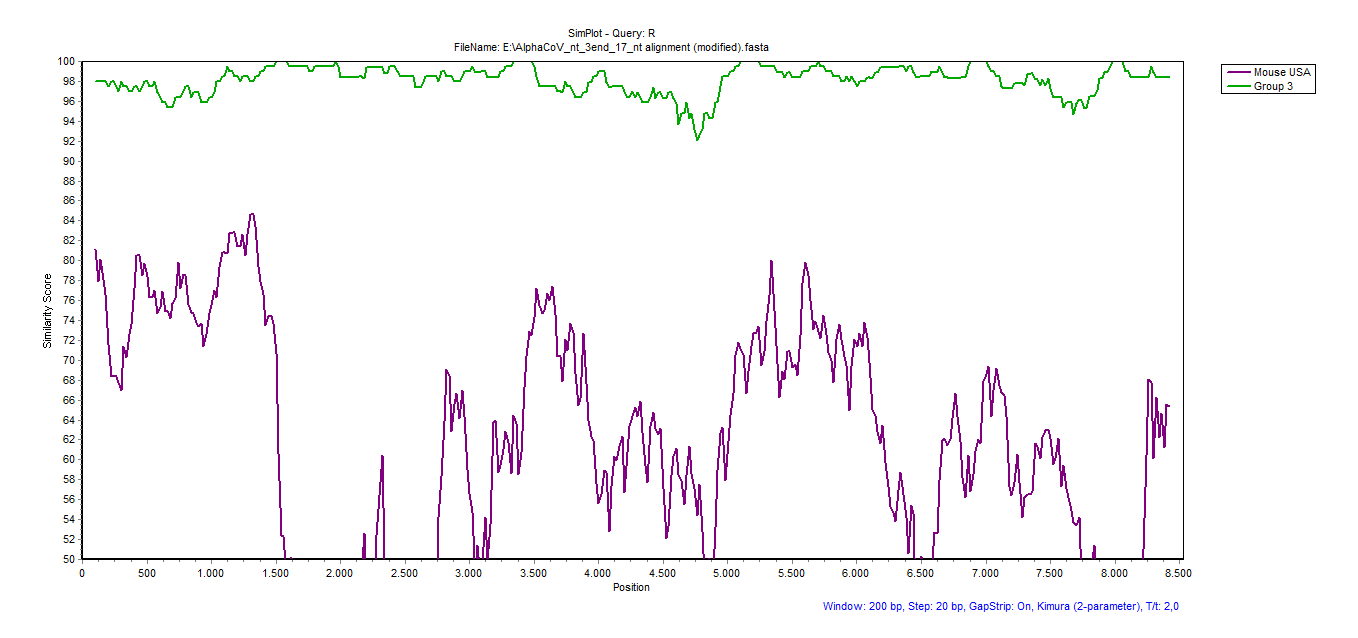

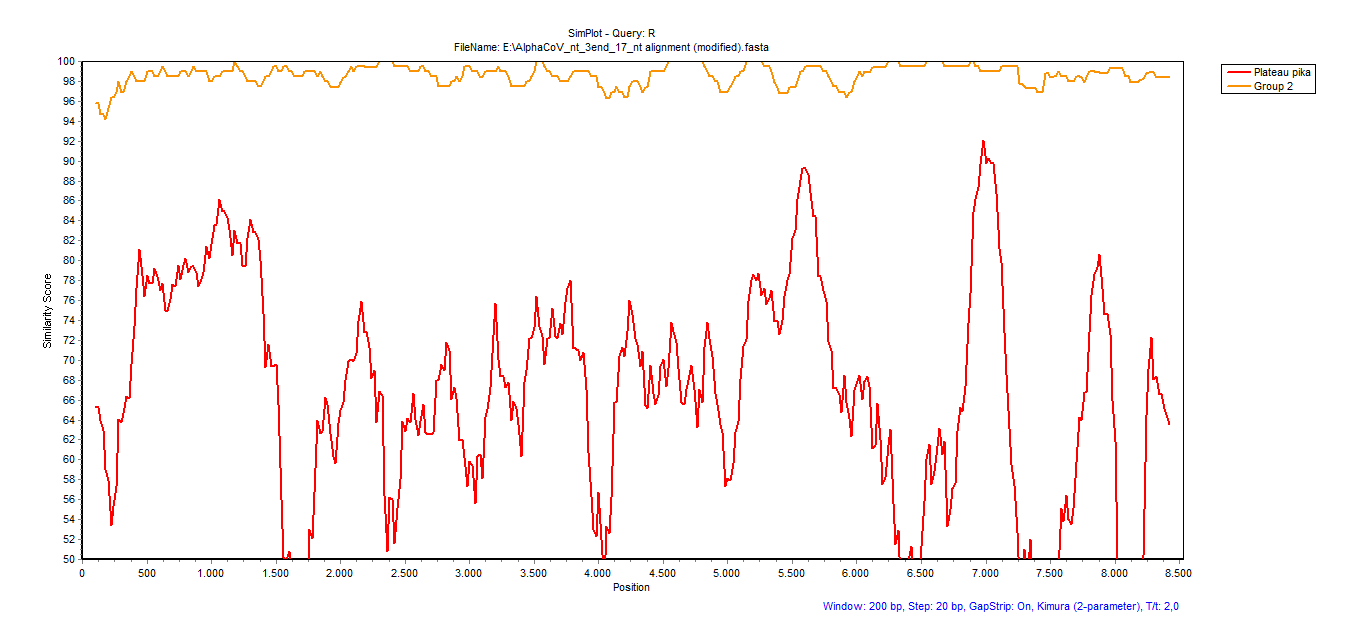
